# Rethinking glottal midline detection

**DOI:** 10.1101/2020.08.20.257428

**Authors:** Andreas M. Kist, Julian Zilker, Pablo Gómez, Anne Schützenberger, Michael Döllinger

## Abstract

A healthy voice is crucial for verbal communication and hence in daily as well as professional life. The basis for a healthy voice are the sound producing vocal folds in the larynx. A hallmark of healthy vocal fold oscillation is the symmetric motion of the left and right vocal fold. Clinically, videoendoscopy is applied to assess the symmetry of the oscillation and evaluated subjectively. High-speed videoendoscopy, an emerging method that allows quantification of the vocal fold oscillation, is more commonly employed in research due to the amount of data and the complex, semi-automatic analysis. In this study, we provide a comprehensive evaluation of methods that detect fully automatically the glottal midline. We use a biophysical model to simulate different vocal fold oscillations, extended the openly available BAGLS dataset using manual annotations, utilized both, simulations and annotated endoscopic images, to train deep neural networks at different stages of the analysis workflow, and compared these to established computer vision algorithms. We found that classical computer vision perform well on detecting the glottal midline in glottis segmentation data, but are outperformed by deep neural networks on this task. We further suggest GlottisNet, a multi-task neural architecture featuring the simultaneous prediction of both, the opening between the vocal folds and the symmetry axis, leading to a huge step forward towards clinical applicability of quantitative, deep learning-assisted laryngeal endoscopy, by fully automating segmentation and midline detection.

## Introduction

Many effective, biological moving sequences are based on symmetric processes, such as walking, running and the generation of voice. The latter is characterized by a passive, symmetric oscillation of the two vocal folds in the larynx (Fig. 1a,b). During healthy phonation, the vocal folds are moving symetrically in regular cycles to the center towards a thought midline where they are touching and then burst outwards (Fig. 1c, Supplementary movie 1). Typical oscillation frequencies are in the range of 80–250 Hz for normal, adult speaking voice (1). If this symmetrical oscillation is impaired, the affected are suffering from a weak, dysfunctional voice (1). Disturbances in symmetry can be caused by organic disorders, such as polyps or nodules. These can be diagnosed easily by normal endoscopy. However, functional disorders are not easily to diagnose properly as there are no visible anatomical changes of the vocal folds. The gold standard of diagnosis, laryngeal stroboscopy, is also not able to catch irregular oscillation patterns due to its underlying stroboscopic principle (2). High-speed videoendoscopy (HSV), a promising tool to quantify the oscillation behavior on a single cycle level, is theoretically able to determine the degree of symmetry of the two vocal fold oscillation patterns (2, 3). The glottal area is typically utilized as a proxy for vocal fold oscillation (4–7) and is extracted from the endoscopy image using various segmentation techniques (e.g. (8–10), as shown in Fig. 1c). To assign a fraction of the glottal area to the left or the right vocal fold, a symmetry axis, or midline, is defined to split the glottal area in two areas (Fig. 1d). This midline detection approach is commonly performed after segmenting the glottal area (11). The glottal area for the individual vocal fold allows a sophisticated analysis of the vocal fold-specific glottal area waveform (GAW), and the phonovibrogram (PVG), a two-dimensional representation of the oscillation behavior and symmetry ((11, 12), see Fig. 1d).

**Fig. 1.**
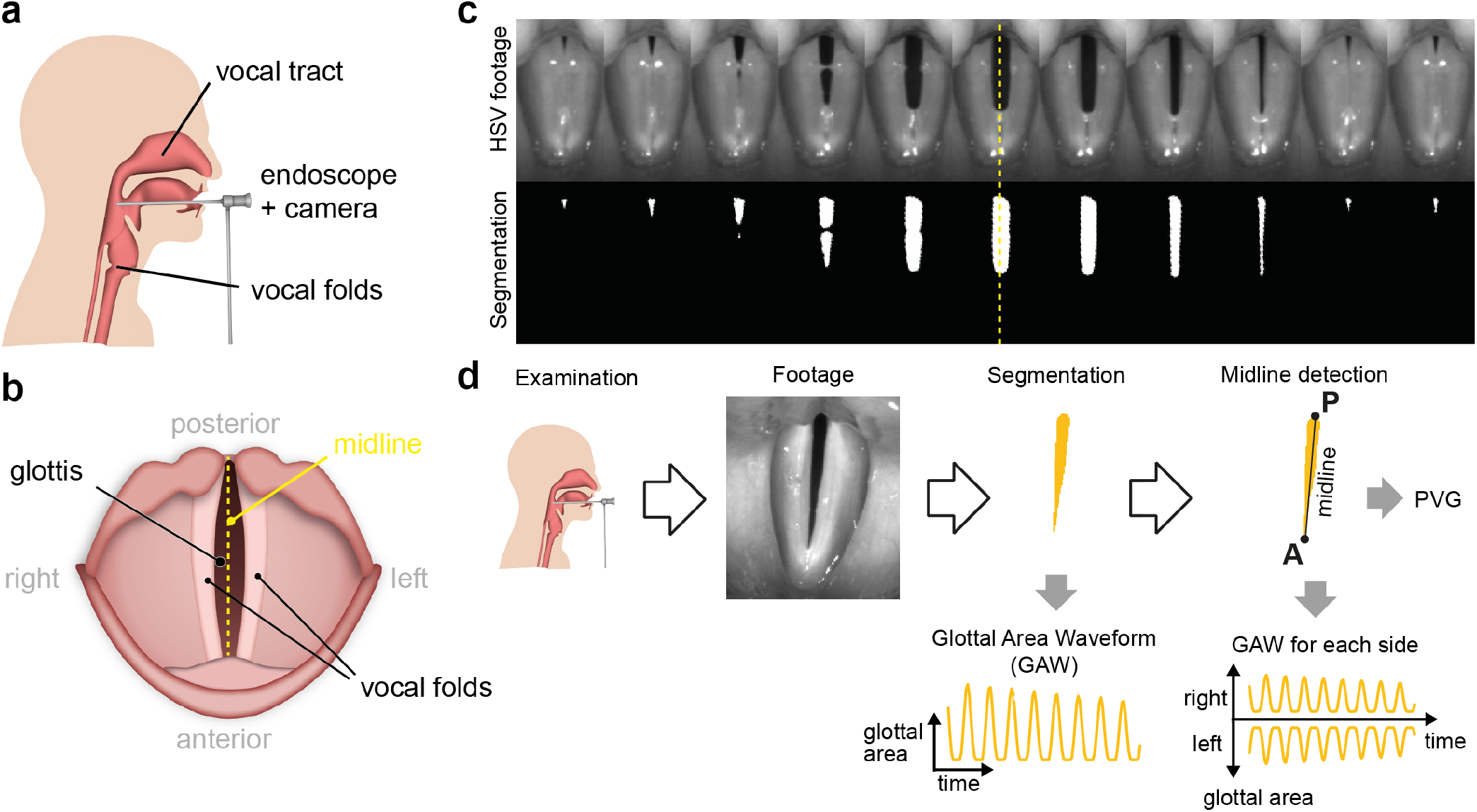
The glottal midline is crucial to compute clinically relevant dynamic left-right symmetry parameters. a) High-speed videoendoscopy examination setup, b) top view onto the vocal folds during high-speed videoendoscopy c) a single HSV oscillation cycle from a healthy individual together with its corresponding glottis segmentation mask. Note the symmetry to the yellow dashed midline. d) State-of-the-art workflow to determine the glottal midline. HSV footage gained from examination is first segmented and converted to a glottal area waveform. On local maxima, the midline is predicted via the posterior (P) and anterior points (A). Using the midline, the GAW for each vocal fold and the phonovibrogram (PVG) can be computed.

As a symmetric vocal fold oscillation pattern is a hallmark of healthy phonation. Previous studies used several methods to define the glottal midline to describe the symmetry. Many works set the glottal midline manually (13, 14), e.g. between the anterior commissure and the arytenoids (13). Recently, the midline was automatically detected using linear regression and interpolation techniques (15) or using principal component analysis (16) to describe vocal fold-specific dynamics. Linear regression, however, potentially underestimates the midline slope, as it by definition only minimizes only residuals in one direction. Phonovibrograms that rely strongly on the midline computation originally use only the top-most and bottom-most point, defined as posterior and anterior point, respectively, identified in the segmentation mask when the glottis is maximally opened (11, 12). Despite the fact that these methods were intensively validated on clinical data, they were only compared to a manual, subjective labeling. A judgement on an objective ground-truth, e.g. using well-defined synthetic data, would be advantageous. Additionally, if the glottis is not completely visible in the footage, both methods are prone to under- or overestimate the glottal midline. As these methods are based on glottis segmentations, the segmentation itself can have huge impact on the midline prediction, and thus, the laryngeal dynamics interpretation.

The symmetric oscillation of the vocal folds have already been described in the first two mass model introduced in (17). Lumped mass models have been shown to be able to model asymmetric oscillation patterns (18–22) and can potentially be used as a source to not only create single mass trajectories as shown before, but also to generate time-variant synthetic segmentation masks. With that, one is able to generate GAWs with by design known properties, such as the glottal midline, and posterior and anterior point. However, no such applications has been reported to our knowledge so far.

In this study, we explore which methods are able to predict the glottal midline accurately in clinical and synthetic data. We further compare which data source, endoscopic images or segmentation masks are the best suited source for midline estimation, and how classical computer vision techniques compare to state-of-the-art deep neural networks. We found that computer vision methods are competitive in predicting the glottal midline in segmentation masks with deep neural networks. Incorporating time information further improves prediction accuracy in both, neural networks and computer vision algorithms. We suggest a novel multitask architecture, that predicts both, glottal midline and glottis segmentation simultaneously in endoscopic footage.

## Results

### A biophysical model creates symmetric and asymmetric time-variant segmentations

An objective performance evaluation of any algorithm or neural network is, for example, a comparison to ground-truth data. As this is not unequivocally possible directly in endoscopic images (see also later paragraphs), we investigated if we can utilize lumped mass models that have previously been shown to accurately model vocal fold oscillation physiology (17, 18, 23) in order to generate high-quality, time-variant segmentation masks. By design, the model defines a midline and thus, a ground truth. We optimized a previously published, established six mass model (6MM, (18)) and simplified it to gain glottal area segmentation masks (Fig. 2a) and GAWs (Fig. 2b). In contrast to the original model described in (18), our adjusted model does not feature negligible movement in the longitudinal direction (24, 25), resolves issues with the damping formulations and has a corrected term for the distance of the masses (see Methods). Using our 6MM implementation, we are able to produce symmetric and asymmetric oscillation patterns (Fig. 2b,c and Supplementary Movie 2) together with translational and rotational motion over time to simulate examination motion artifacts. Additionally, we introduced noisy pixels to the segmentation mask contour to generate segmentation uncertainty. We created a synthetic evaluation dataset featuring 2500 simulations with by design known gottal midline allowing to objectively judge the performance of methods that use segmentation masks to predict the glottal midline.

**Fig. 2.**
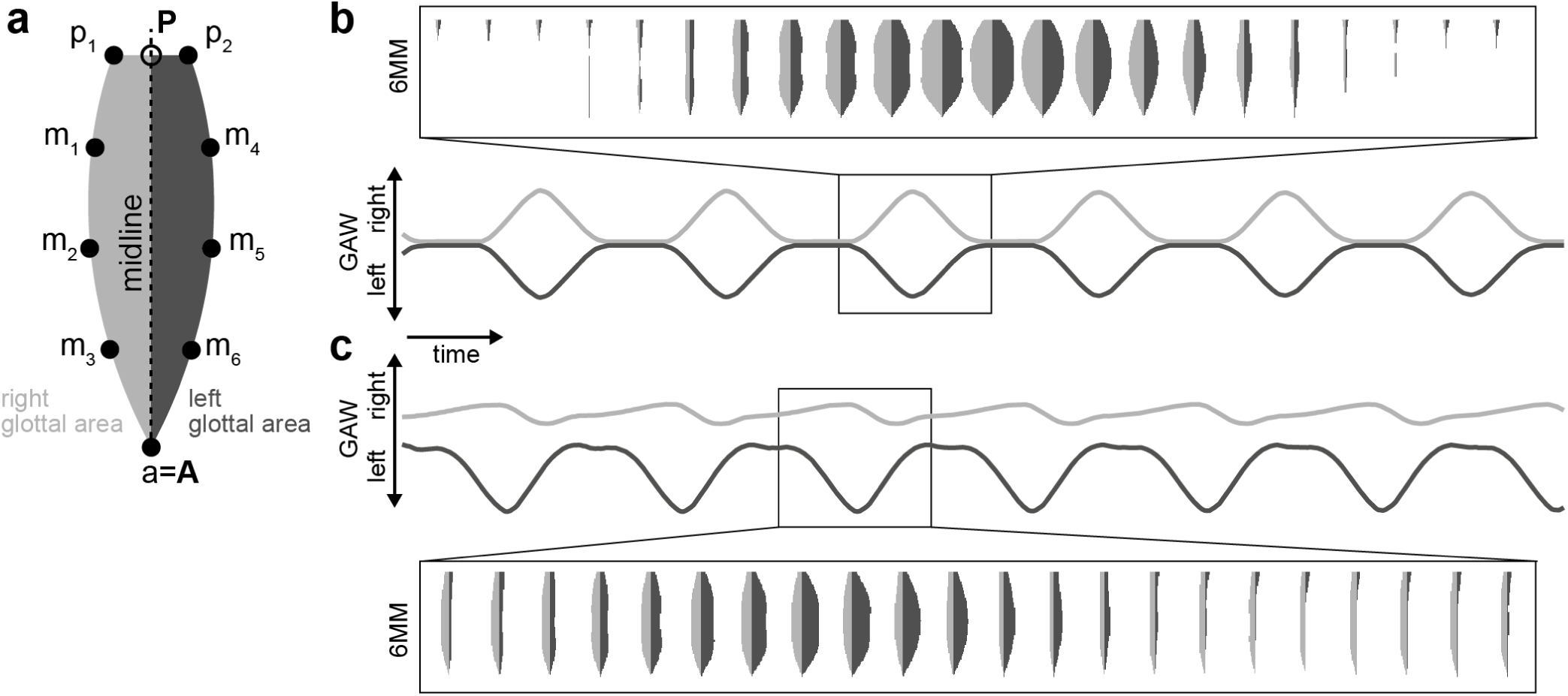
The six mass model (6MM). a) an example glottal area, split into left (dark gray) and right (light gray) using the glottal midline connecting posterior (**P**) and anterior (**A**) point. The six movable masses (m1-m6) are arranged left and right from the glottal midline. The posterior point can be divided into two fixed masses (p1 and p2) inducing a posterior gap, as seen in healthy female individuals (see also panel b), whereas the anterior point (a) is a fixed mass. b) Symmetric oscillation of the left and right masses result in symmetric glottal area waveforms (GAWs) indicated in the same gray color as shown in panel a). Glottal area model output is shown for an example cycle. c) same arrangement as in b), however, with an asymmetric oscillation pattern. Note the right vocal fold insufficiency leading to an always partially open glottis on the right side.

To test algorithms on *in vivo* data, we used the Benchmark for Automatic Glottis Segmentation (BAGLS) dataset (10). We extended BAGLS by manually annotating the training and the test dataset of single endoscopic images using anatomical landmarks, such as anterior commissure and arytenoid cartilages (see Methods).

### Computer vision methods accurately predict midline in symmetric oscillations

In previous studies, the anterior and posterior point is predicted from the segmentation masks in each opening-closing cycle, where the glottis is maximally opened (Fig. 3a), e.g. (12). Therefore, we first focused on segmentation mask-based midline prediction methods following this approach. We evaluated classical computer vision methods in this study, which do not rely on previous (learned) knowledge in contrast to popular deep neural networks. We found that orthogonal distance regression (ODR), principal component analysis (PCA), Image Moments and Ellipse Fitting are potentially able to predict the glottal midline (table 3 and Methods) and compared them to previously described methods, i.e. top most and bottom most points (TB) and linear regression (LR) as described in (15) and (12), respectively. An overview of the computer vision algorithms and their working principle is shown in Supplementary Fig. 1 and explicitly described in the Methods. Briefly, ODR fits a linear equation by minimizing residuals in both dimensions, in contrast to LR, where only residuals in one dimension are minimized. PCA is converting the image space to a principal component space, where the first principal component is along the highest variance and thus, approximates the midline. Moments are a common principle in mathematics and used in fields such as classical mechanics and statistical theory, but have been applied intensively to the image processing field since the 1960s (26, 27), and describe the midline using the center of mass of the image and a direction vector. Last, by fitting an ellipse to the glottal area outline, the major axis can be interpreted as equivalent to the midline.

**Table 1.** Comparison of different neural network architectures to predict glottal mid-line in segmentation masks. *We also tested NasNet-Large, but it did not converge.

| neural network | median mIoU | parameters |
| --- | --- | --- |
| VGG19 | 0.691 | 20.0M |
| NasNet-Mobile* | 0.804 | 4.2M |
| InceptionV3 | 0.812 | 21.8M |
| *MidlineNet* | 0.874 | 0.1M |
| ResNet-50 | 0.878 | 23.5M |
| MobileNetV2 | 0.884 | 2.2M |
| U-Net encoder | 0.931 | 9.4M |
| EfficientNetB0 | 0.936 | 4.0M |
| Xception | **0.960** | 20.8M |

**Table 2.** Overview of midline estimation procedures and their respective advantages and disadvantages

| Source | Method | Advantage | Disadvantage |
| --- | --- | --- | --- |
| Endoscopy image | Deep neural network | Direct midline estimation, not restricted to segmentation | Subjectivity in anatomical landmarks |
| Segmentation | Computer vision alg. | Universal and fast | Issues with asymmetric conditions, underestimate glottis length |
|  | Deep neural network | Can be trained on synthetic data |  |

**Table 3.** Overview of classical computer vision algorithms and their respective midline descriptor together with references where the respective algorithm was applied to glottal midline detection. *(16) also uses PCA combined with custom optimization strategies.

| Method | Reference | Midline descriptor |
| --- | --- | --- |
| Top and bottom most point (TB) | (15) | Line connecting top and bottom most point in y |
| Linear regression (LR) | (12) | Optimization of linear function by minimizing residuals in y |
| Orthogonal Distance Regression (ODR) | *this paper* | Optimization of linear function by minimizing residuals in x and y |
| Principal Component Analysis (PCA) | *this paper\** | 1st principal component |
| Image Moments | *this paper* | Center of mass and orientation vector |
| Ellipse Fitting | *this paper* | Major axis |

**Fig. 3.**
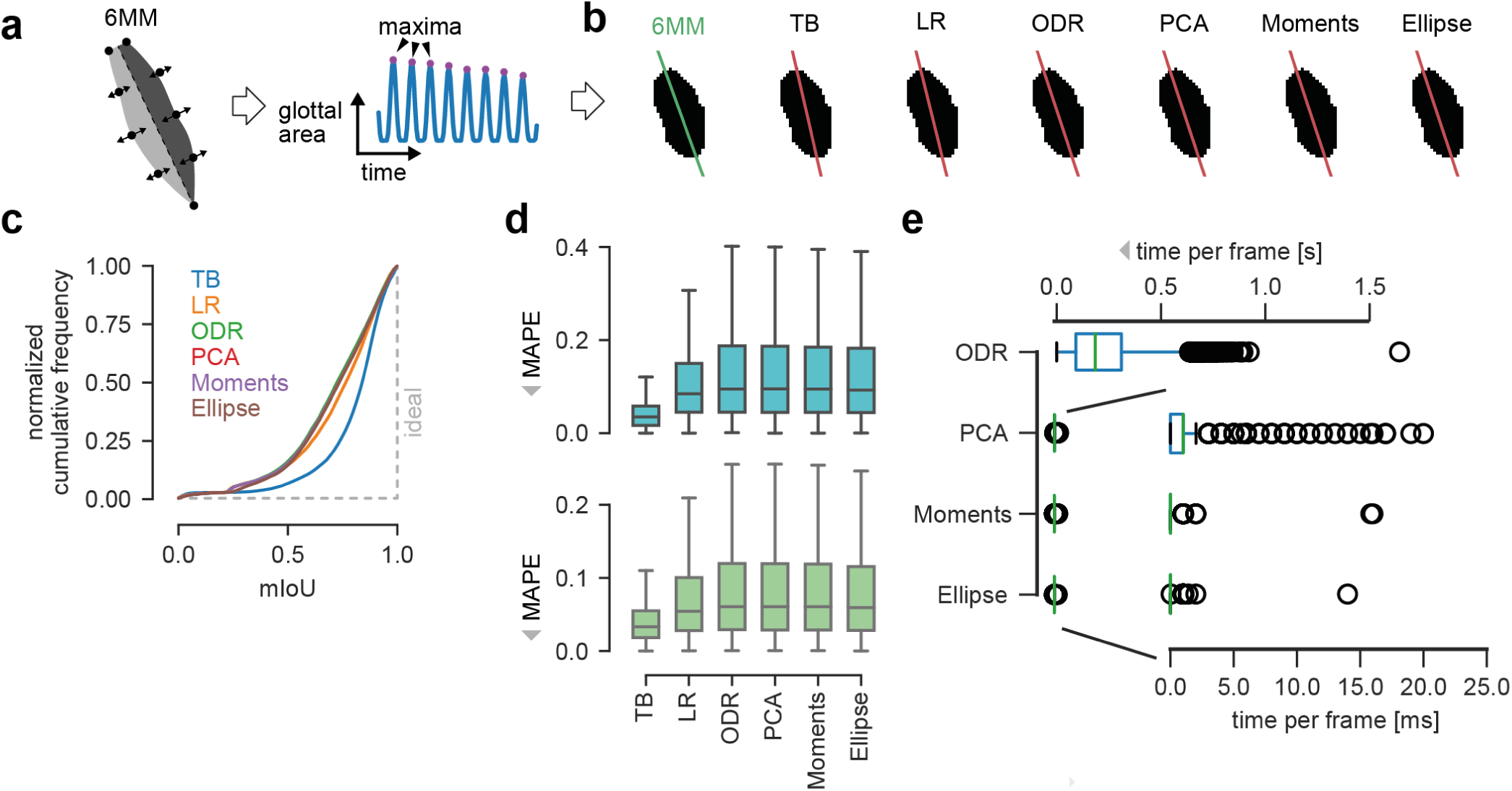
Performance of computer vision algorithms. a) Evaluation procedure. GAWs are computed from 6MM simulations. Maxima were found and respective image frames were analyzed by computer vision algorithms. b) Midline predictions of computer vision algorithms (red lines) compared to ground-truth (green) on exemplary 6MM data. c) Cumulative mIoU scores across the synthetic dataset for all algorithms tested. The ideal curve is indicated as gray dashed line. d) Distribution of the mean absolute percentage error (MAPE) for posterior (cyan) and anterior (green) point across the synthetic dataset for all algorithms tested. e) Computation time of new algorithms tested. TB and LR required virtually no computation time.

We analyzed the performance of the various algorithms in terms of midline prediction accuracy and speed. We measured accuracy in two ways: first, the relative euclidean distance, as measured by the mean absolute percentage error (MAPE), between prediction and the ground-truth posterior and anterior point, and second, the mean intersection over union (mIoU) for the left and right glottal area using the prediction and the corresponding ground truth (see Methods and Supplementary Fig. 2). The use of both metrics is particularly important, because small MAPE scores could have a tremendous effect on the resulting glottal areas, whereas points that are moved along the midline can show large MAPE scores, but still result in highly overlapping glottal areas with the ground-truth.

We first tested all algorithms on a toy dataset consisting of an ellipse as rough approximation of a segmented glottis rotated by a defined angle to ensure correct implementation of each algorithm and to determine the valid range of rotation angles (Supplementary Fig. 3, Supplementary movie 3). Our data suggests that TB and LR are prone to under- or overestimate the midline of a perfect symmetric object, and that ODR, PCA, Image Moments and Ellipse Fitting perfectly identify the midline with almost zero relative distance (Supplementary Fig. 3b,c,d). Further, using this toy dataset, the mIoU score for ODR, PCA, Image Moments and Ellipse Fitting is always close to 1, whereas the mIoU score for TB and LR varies tremendously (Supplementary Fig. 3e). Especially rotation angles beyond ± 30° cause severe artifacts in TB and LR (Supplementary Fig. 3b,c,e). The BAGLS dataset that contains representative clinical data shows that typical rotation angles as determined by rectifying the glottis using PCA are ranging from −30 to +30 degrees (Supplementary Fig. 4) and appear to be Gaussian distributed (6.98 ± 10.1 degrees). The non-zero mean may be caused by the high prevalence of right-handed examiners. Taken together, ee include TB and LR in further analyses and apply a random rotation of [-30°, 30°] to our synthetic dataset for data augmentation purposes and mimic a realistic clinical setting.

We calculated the GAWs from all 2500 simulations that were generated by our 6MM and detected the local maxima in each GAW (Fig. 3a). For each local maximum, the glottal midline was predicted by each algorithm from the corresponding segmentation mask (Fig. 3b). We found that all algorithms are able to predict an accurate glottal midline in symmetric cases as exemplary shown in Fig. 3b. However, especially in the important asymmetric cases, these algorithms have problems in properly predicting the glottal midline (Supplementary Fig. 5), leading to low mIoU scores (Fig. 3c). Interestingly, methods that are not equally accounting x and y, i.e. TB and LR, are superior to other methods, such as ODR, PCA, Image Moments and Ellipse Fitting (Fig. 3c). A similar trend is visible in the MAPE scores (Fig. 3d), where TB has lower scores, whereas the distributions of the other methods are not differing from each other.

The computation time across algorithms is heterogeneous (Fig. 3e). The TB and the LR algorithms are computed almost instantaneously (*<* 1 ms) and are therefore not represented in the Figure. PCA, Image Moments and Ellipse Fitting are also highly efficient and their computation in most cases takes less than 1 ms. On roughly 1% of the images, however, PCA took longer than the very efficient Image Moments and Ellipse Fitting routines (2 ms and 1 ms) with a maximum duration of 20 ms compared to 14 ms and 16 ms for Image Moments and Ellipse Fitting, respectively. However, ODR was significantly slower and the algorithm took 229 ms on average to finish, but 7% of the images took more than 500 ms to converge (Fig. 3c).

In summary, all methods are able to predict an accurate midline in many cases, however, TB is outperforming the others in terms of accuracy (Fig. 4c,d) and computational efficiency. ODR has similar performance as PCA, Image Moments and Ellipse Fitting, however, needs way longer to be computed, suggesting that ODR is inferior.

**Fig. 4.**
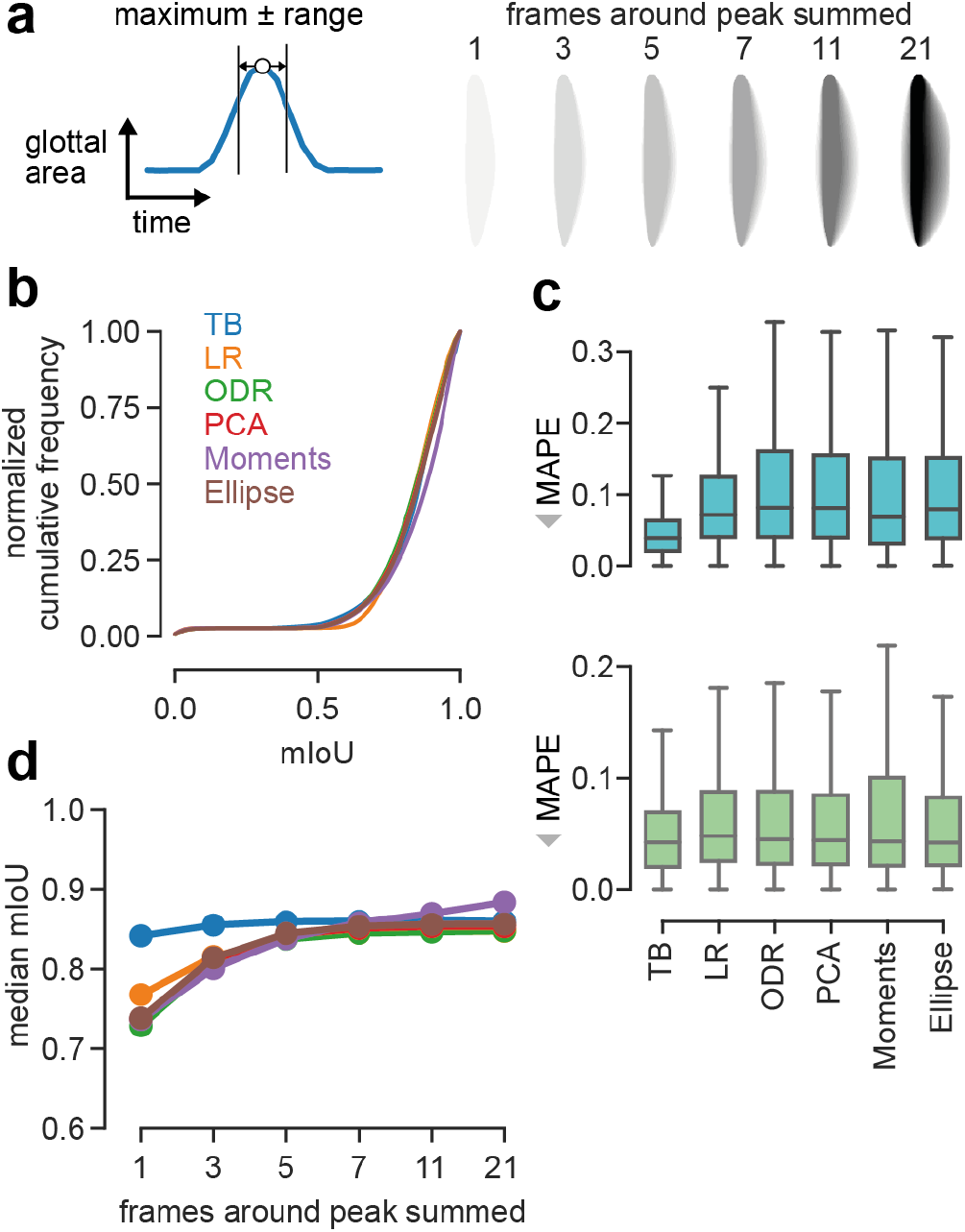
Introducing time increases performance in most algorithms. a) Maximum is detected in GAW. The respective frame together with a pre-defined range is summed over time. An example of summing multiple frames over different ranges are shown on the right. b) Cumulative mIoU scores for algorithms tested when con-sidering a total of 21 frames. c) MAPE scores of algorithms for posterior (cyan) and anterior (green) points when considering 21 frames. d) Median mIoU scores for dif-ferent algorithms compared to the amount of frames summed around the detected maximum peak. Same color scheme as in b).

### Providing temporal context increases performance of computer vision algorithms

The established approach of using only the segmentation mask when the glottis is maximally opened, has the downside that the temporal context and hence, important oscillation behavior is lost. Together with the underlying principles of PCA, Image Moments, Ellipse Fitting and ODR, a symmetric distribution of the data is assumed. We therefore evaluated the computer vision algorithms on images that not only contain the segmentation mask of the maximum opened glottis, but also prior and succeeding frames (Fig. 4a, left panel). By using different ranges, the real midline, i.e. oscillation center, is also visually more apparent and easier to determine compared to a single frame (compare different ranges in Fig. 4a, right panel). When using a range of up to 21 frames (ten frames on each side of the peak and the frame containing the peak), the cumulative mIoU scores improved across all methods, except TB (Fig. 4b). Interestingly, Image Moments is slightly better than the other methods in terms of median mIoU scores when considering a total of 7, 11 and 21 frames (Fig. 4b, Fig. 4d). All methods are further able to accurately predict the anterior point (Fig. 4c, median MAPE = 0.042, 0.048, 0.045, 0.044, 0.043 and 0.042 for TB, LR, ODR, PCA, Image Moments and Ellipse Fitting, respectively), in contrast to the single frame prediction (Fig. 3d). However, the posterior point is still best predicted by the TB method (median MAPE=0.039), and worse, but similar predicted by the other methods (median MAPE=0.072, 0.082, 0.082, 0.069 and 0.0.080 for LR, ODR, PCA, Image Moments and Ellipse Fitting, respectively).

In summary, a single frame is sufficient for the TB method to outperform other algorithms in terms of mIoU. However, by providing temporal context, i.e. more frames (*>* 7), to the algorithms, they reach similar performance (LR, ODR, PCA, Ellipse Fitting) or are even outperforming the TB method (Image Moments).

### Deep neural networks outperform classical methods on asymmetric oscillations

We next investigated how deep neural networks compare to classical methods. We evaluated several state-of-the-art architectures that have been used for feature extraction and are established in the field. In particular, we evaluated the ResNet (28), Xception (29), InceptionV3 (30), NasNet (31), EfficientNet (32), MobileNetV2 (33) and VGG19 (34) architectures (overview table 1. Additionally, we tested the U-Net encoder (35). The rationale for using the U-Net encoder is that the U-Net itself has been shown to perform well on glottis segmentation tasks (36). We found that we could optimize the filter structure in the U-Net encoder to gain a lean and fast, yet high performing model (*MidlineNet*, see Methods) that we additionally evaluate in this study.

We first tested how neural networks perform on the same two tasks (single or multiple frames combined in one image, Fig. 5a, upper panel). Using only the frame of the GAW maximum, we found that there are apparent performance differences across architectures (overview of all architectures in Supplementary Fig. 6). Consistently, the VGG19 architecture resulted in the worst median mIoU score (0.691, Fig. 5b and table 1), followed by NasNet-Mobile (0.804) and InceptionV3 (0.812). As shown in Fig. 5b,c and table 1, MidlineNet, MobileNetV2, U-Net encoder, EfficientNetB0 and the Xception architecture are outperforming classical computer vision methods operating on a single frame. The ResNet-50 architecture is also outperforming computer vision methods, however, due to the large parameter space and similar performance as the MidlineNet architecture (Fig. 5h), we decided not to follow up on this architecture. We found similar results for the MAPE metric, where almost all architectures expect VGG19 and NasNet-Mobile outperformed the best computer vision method (Supplementary Fig. 7a,b). We next investigated if the top performing neural networks further increase their performance when also providing adjacent frames similar to Fig. 4. Indeed, when trained on multiple frames overlaid, networks tend to show superior performance (Fig. 5d,h). For some configurations, we observe small performance drops, for example for 7 or 21 frames overlaid in EfficientNet, that is maybe due to variations for individual seeds. Especially the MidlineNet architecture benefits from the additional time information, achieving almost the same performance as more sophisticated architectures: the mIoU score for 21 overlaid frames is 0.941, which is an increase of 7.6%. We show that the Xception architecture consistently outperforms other architectures (best mIoU = 0.974), indicating that the Xception architecture utilizes its higher parameter space. The already low MAPE scores (*<* 0.1 for a single frame) were slightly lowered in some cases (Xception and MidlineNet), in other cases, the temporal context did not consistently improve the MAPE score (Supplementary Fig. 7c,d), indicating that low MAPE scores alone do not fully reflect the midline prediction accuracy.

**Fig. 5.**
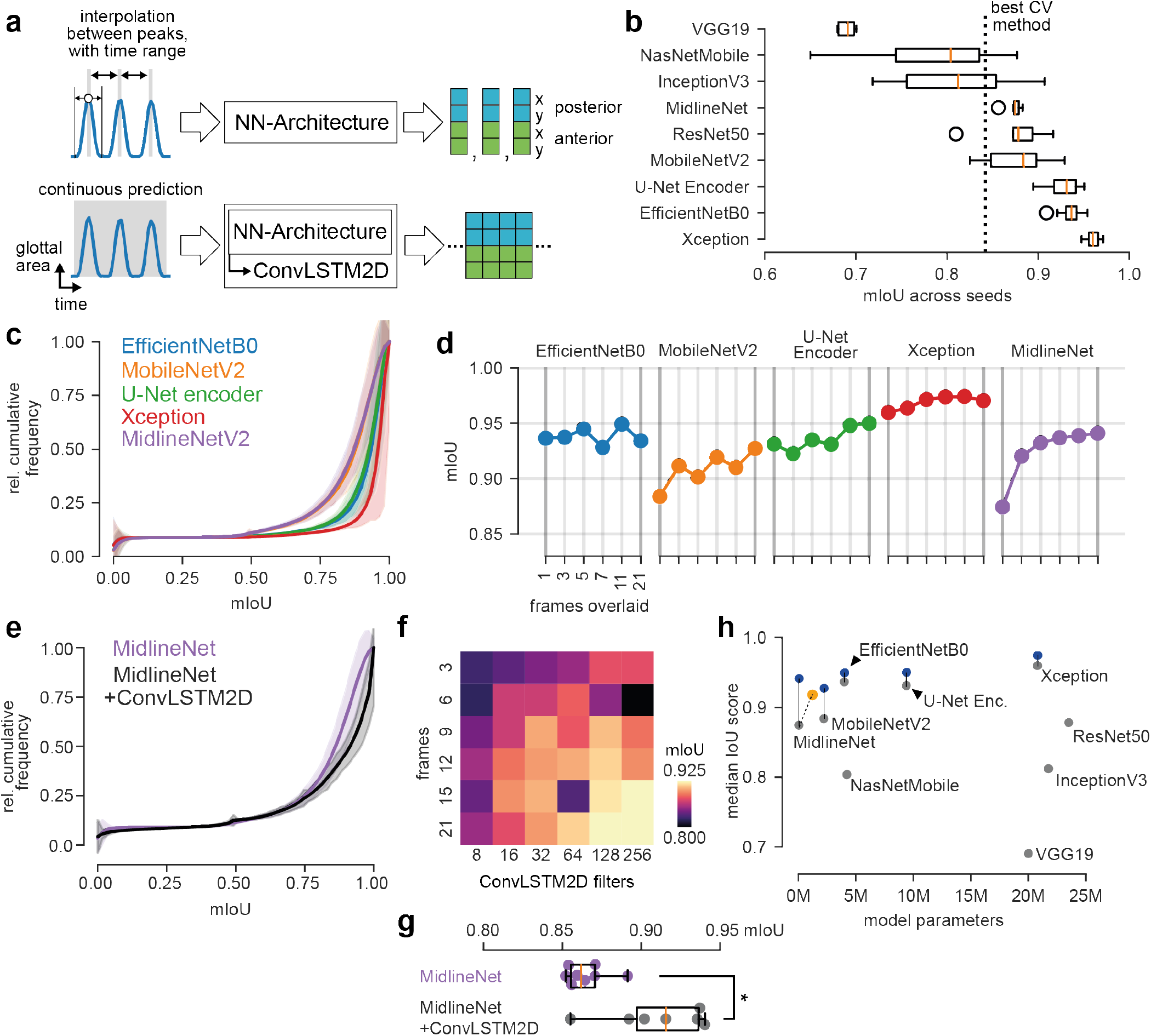
Deep neural networks outperform classical computer vision methods. a) Overview of prediction methods. Either a neural network architecture directly predicts anterior and posterior point coordinates from the maximum opened glottis (and a range of adjacent frames, optionally), or it uses a history-based approach together with LSTM cells to predict continuously posterior and anterior point coordinates. b) Distribution of median mIoU scores across different neural network architectures and seeds on the test set. c) Average cumulative mIoU scores on the test set for selected neural architectures shown in d). Shaded error indicates standard deviation. d) Median mIoU scores for different neural architectures and varying temporal context. e) Average cumulative mIoU scores of the U-Net encoder and its ConvLSTM-variant. Shaded error indicates standard deviation. f) Colormap of median mIoU scores depending on sequence length and ConvLSTM2D filters. g) Distribution of median mIoU scores U-Net encoder and its ConvLSTM-variant across different seeds. h) Overview of neural network performance depending on size. Gray circles indicate baseline performance (single frame inference) and blue circles indicate temporal context by summing frames. Yellow circle indicates the MidlineNet ConvLSTM-variant.

For time-series forecasting, recurrent neural networks are typically applied (37). In a recurrent setting, single frames for each time step are provided and analyzed, in contrast to the aforementioned approach where frames were summed over time, losing the frame-by-frame resolution. We use our MidlineNet as an example neural architecture to showcase the effect of ConvLSTM2D layers. Here, it is straightforward to adapt MidlineNet to a recurrent convolutional neural network by changing the Conv2D layers to ConvLSTM2D layers, in contrast to other architectures studied in this context. ConvLSTM2D layers are recurrent layers that internally use convolutions, important for processing 2D data, such as images, instead of matrix multiplications (38). We first performed a hyperparameter search investigating appropriate settings for ConvLSTM2D filters and frames fed to the network. We found that 15 to 21 frames together with 128 to 256 filters provided the best performance with an median mIoU of greather than 0.9 (Fig. 5f). When comparing the ConvLSTM2D-variant to its static counterpart, we found that the recurrent variant performs significantly better (Fig. 5g, p<0.05, Student’s t-test), indicating that further research using ConvLSTM2D-based architectures is important. However, directly integrating the time information into the image by overlaying the frames shows even higher performance (Fig. 5d,g,h).

In summary, we found that deep neural networks have varying performance on midline prediction, where the Xception architecture consistently achieved good results (Fig. 5b,c,d). We introduced time in two different ways, i.e. overlaying multiple frames and provide the merge to a convolutional neural network, and providing single, adjacent frames to a recurrent neural network. We found that both ways have improved performance compared to the single static image. Despite the fact that recurrent neural networks are able to continuously predict the midline, we find that summing prior and succeeding frames is easier to implement, allows to use a higher variety of architectures, and yields even greater performance.

### GlottisNet predicts midline and segmentation mask simultaneously

Similar to earlier studies (11, 15), we focused on detecting the glottal midline based on the glottal area segmentation that is typically derived from endoscopic images (sequential path, Fig. 6a). However, other studies already estimated the midline manually in endoscopic images (13, 14) to ensure an unbiased view given the anatomical environment. This also has the advantage that anterior and posterior point are defined as real anatomical landmarks and not as upper and lower intersection of the glottal midline with the segmentation mask. We therefore investigated the fully automatic prediction of the glottal midline directly in endoscopic images to avoid a prior segmentation step. To answer this question, we utilized the openly available BAGLS dataset (36), and annotated manually the anatomical anterior and posterior point in all 59,250 images (Supplementary Fig. 8), and thus, defining the glottal midline. We focused on encoder-decoder architectures, as these have been shown to reliably segment the glottal area (9, 36), and integrated the anterior and posterior point prediction into the architecture. As baseline we use the U-Net architecture ((35), Fig. 6b). With that, we introduce a novel multitask architecture, that simultaneously provides both, glottal area segmentation and anterior/posterior point prediction (simultaneous path, Fig. 6a,b).

**Fig. 6.**
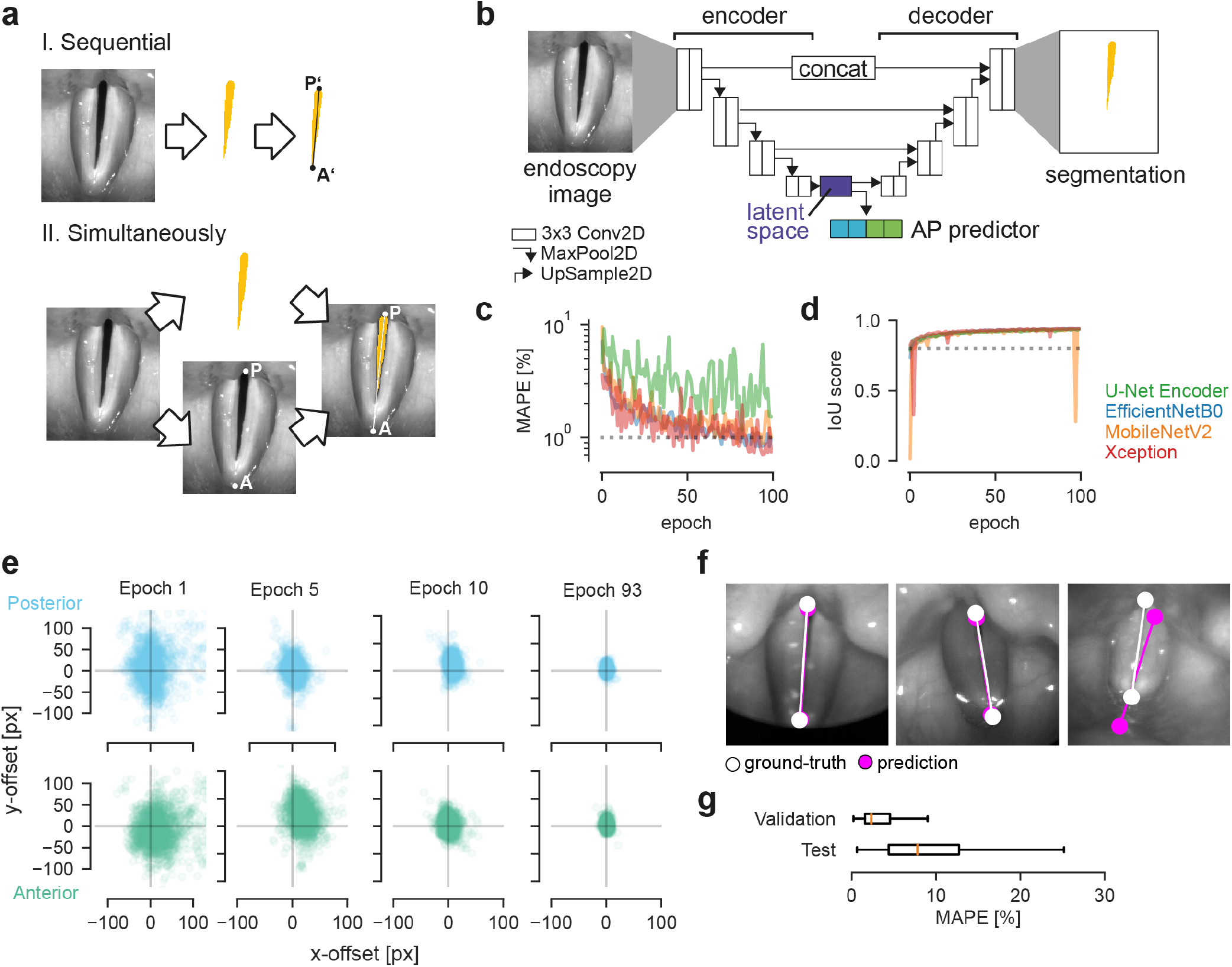
GlottisNet is a multi-task architecture that predicts simultaneously both, glottal midline and area. a) Comparison of sequential (upper panel) and simultaneous prediction (lower panel) of the glottal midline. Note the differences of P and A points. b) General GlottisNet architecture consisting of an encoder-decoder network with an additional AP predictor. c) Convergence of MAPE across training epochs for different encoding backbones. d) Convergence of IoU score across training epochs for different encoding backbones. e) X-Y accuracy of P and A point prediction. f) Example images of endoscopic images with glottal midline ground-truth and prediction. g) Distribution of MAPE scores across validation and test dataset for GlottisNet with Xception backbone.

#### The latent space is best suited for midline prediction

Because endoscopic images feature higher variability than binary segmentation masks, we first evaluated which loss is suitable for training. For a preliminary evaluation, we chose the U-Net encoder backbone. We found that any of the tested losses, i.e. mean average error (MAE), mean squared error (MSE), Huber loss and Log-Cosh loss (see Methods), are able to train the network, however, MAE and MSE consistently yielded the best score on the validation data without any sign of divergence (Supplementary Fig. 9). We thus decided to use the MSE loss in any further training procedure. We further found that network convergence in terms of key-point prediction is best when using the latent space or layers in the decoder as input (Supplementary Fig. 10). We found that the multitask optimization does not negatively affect the segmentation performance (Supplementary Fig. 10). For further experiments, we decided on using the latent space as input for A and P point prediction and as entry point for the segmentation decoder (Fig. 6b).

#### GlottisNet is a powerful multitask architecture

As the U-Net is based on an encoding-decoding network (Fig. 6b), we tested if changing the encoder backbone can yield improvements in performance. In particular, we evaluated the vanilla U-Net encoder architecture (35), together with the high performing networks from Fig. 5, namely the MobileNetV2, the EfficientNetB0, and the Xception architecture. We found that all encoder combinations yielded similar, high IoU scores for the segmentation task, but varied largely in their performance predicting A and P point (Fig. 6c,d). Interestingly, all architectures have very similar performance on training and validation set for the segmentation task. However, in terms of A and P point prediction, the U-Net encoder is consistently worse in both, training and validation (Supplementary Fig. 11). Further, the segmentation task is easily learned after only a few epochs (Fig. 6d), the A and P point prediction task takes around 100 epochs to converge to satisfying performance levels (Fig. 6c, Supplementary Fig. 11). In general, we found that the EfficientNetB0 backbone provides a good converging behavior (as shown as converging, concentric point clouds over time, Fig. 6e and Supplementary Fig. 12a), whereas the other backbones show distinct translational displacement towards the center (Supplementary Fig. 12b-d). We tested all four networks on the BAGLS test dataset and found that the EfficientNetB0 backbone has the best performance (median MAPE = 7.79, mIoU = 0.746), followed by the Xception architecture (9.26, 0.778), MobileNetV2 architecture (13.56, 0.751) and the vanilla U-Net encoder (14.84, 0.777). Our results indicate that the vanilla U-Net encoder is performing well on the segmentation task (similar level as the sophisticated Xception architecture). However, it performs poorly on regression tasks. In contrast, the EfficientNetB0 architecture has a good performance on both tasks. As recently shown (39), all test IoU scores are of sufficient quality. Our GlottisNet architecture with the EfficientNetB0 backbone shows visually reasonable prediction behavior on most images and videos (Fig. 6f, Supplementary Movie 3), and despite the relative large deviation in the MAPE score on the test dataset (Fig. 6g), suggesting that both, visually inspection and quantitative metric should be taken into account.

Overall, we found that using the EfficientNetB0 architecture as backbone for the GlottisNet architecture, we achieve best performance in A and P point prediction and very good performance in glottal area segmentation. This combination allows the successful, simultaneous prediction of both, glottal midline and glottis segmentation, on endoscopic images.

## Discussion

In this study, we provide a comprehensive analysis of methods to estimate the glottal midline from either endoscopic images or segmentation masks. For endoscopic images, we suggest a novel multitask architecture named GlottisNet that allows the simultaneous prediction of both, glottal midline and segmentation mask. We show that both, classical algorithms and state-of-the-art deep neural networks are able to predict accurate glottal midlines on segmentation masks. Further, we show that our modified six mass model (6MM) is a valid tool to generate synthetic, yet realistic, time-variant segmentation masks, needed for an objective evaluation of segmentation-based algorithms. We were further able to show that adding the oscillation history improves the performance significantly (Fig. 4, Fig. 5), similar to previous reports (9).

### Definition of glottal midline

As symmetry is a hallmark for healthy oscillation behavior (1), the definition of the symmetry axis, i.e. the glottal midline, is of great importance for an accurate diagnosis. An anatomical derivation is the most appropriate, yet hard to unequivocally define (13, 14) and maybe a source for interannotator variability (40). Defining the glottal midline from a simpler representation, i.e. glottal area segmentations, is easier and potentially more robust (compare Fig. 3, Fig. 4 and Fig. 5 to Fig. 6), than directly predicting the glottal midline in endoscopy images. However, since recently it has been very challenging to even find these anatomical landmarks automatically in endoscopy footage due to technical limitations and limited availability of labeled data. In our study, we overcome these limitations by extending an open available dataset (36) with manual annotations and combining it with state-of-the-art deep neural networks Fig. 6 to yield GlottisNet. However, our approach is based on single frames and lacks the oscillation behavior to potentially further improve the prediction accuracy (as suggested in Fig. 4 and Fig. 5). Extending GlottisNet into a recurrent neural network is not trivial, we therefore suggest a recurrent version of GlottisNet for a subsequent study.

### Synthetic data to train deep neural networks

Supervised training of deep neural networks is highly dependent on large, annotated data (41). Acquiring and annotating the data is time-consuming and expensive. Using synthetic models that can closely mimic real data is therefore cost-effective and flexible. With our six mass model, we not only produce high quality segmentation masks that closely resemble segmentations derived from real endoscopic footage, but also by design know the oscillation center, i.e. the glottal midline, resembling the perfect ground truth. By using manually annotated data, we would introduce an annotator’s bias, that we circumvent with our strategy. By having ultimate control over the model parameters, we are able to identify strengths and weaknesses of the algorithm, such as performance in varying environments (Supplementary Fig. 3, 5 and Fig. 3, Fig. 4). Already with six moving masses and appropriate post-processing, 6MM-based glottis segmentations are producing exceptional results. However, by using more moving masses and different post-processing steps, one can potentially further improve the synthetic data quality. Additionally, we envision that other ways of generating synthetic data, by using Generative Adversial Networks (GANs, (42, 43)) or Bayesian variational autoencoders (VAEs, (44)) maybe suitable for generating synthetic data.

### Classical computer vision algorithms are worse, but not bad

Since the advent of deep learning methods, classical computer vision algorithms seem to be succeeded by neural networks in several computer vision tasks (41, 45). However, we find that classical computer vision algorithm indeed perform well on our task (Fig. 3, Fig. 4). Notably, these methods can be implemented straight without the need of labeled data or any training procedure. Computer vision algorithms are highly general and show a predictable behavior, whereas the behavior of neural networks on unknown data that may lay outside the training data distribution is rather unpredictable. Interestingly, on a single image the simple heuristic TB performs better than VGG19, NasNet-Mobile and InceptionV3, and across several images all computer vision methods outperform these deep neural networks (compare Fig. 4, Fig. 5 and table 1).

### Clinical impact

Quantitative, symmetry axis prediction dependent parameters, have an impact on diagnosis and treatment options (46–48). Especially clinically relevant phonovibrograms are dependent on the correct detection of the midline (11). As phonovibrograms are based on the extent of the segmentation, they are highly biased towards the maximum opened glottal area, neglecting the total extent of the vocal folds. Incorporating the real anatomical conditions to phonovibrograms via the glottal midline prediction in endoscopic images (e.g. via GlottisNet, Fig. 6), a normalized phonovibrogram would allow better comparability across subjects and may positively influence phonovibrogram-derived disease classifications and quantitative parameters.

In many areas deep neural networks have shown their usability in guiding clinicians and diagnosing diseases and the great potential of artificial intelligence (49–51). With our study, we provide a comprehensive, high-quality toolbox to allow a fully automatic detection of the glottal midline essential for determining clinical relevant parameters. By combining the glottal midline detection together with the glottis segmentation, we overcome the main bottlenecks of clinical applicability of HSV and hence are able to bring quantitative HSV a huge step closer towards clinical routine.

## Methods

In this study, we aim to find the best line that splits the glottal area in two areas representing the oscillation behavior of the left and the right vocal folds. We define the line using a linear equation (Eq. S (1)) that connects both, posterior and anterior point. As some methods are predicting first the slope and intercept, the posterior point and anterior point are defined as the first and last intersection between line and segmentation shape, respectively.

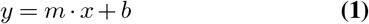

### Biophysical model

Our biophysical model consists of two fixed (p and a), and six moving masses (*m*_1_ to *m*_6_) as shown in Fig. 2a. In women, a small gap at the P point is physiological. We acknowledge this by choosing a random offset between p_1_ and p_2_. The moving mass positions, and thus, the vocal fold dynamics, are described through a system of ordinary differential equations and are derived from (18):

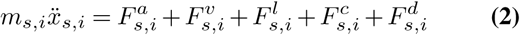

where 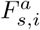 is the anchor spring force, 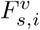 the vertical coupling force, 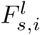 the longitudinal coupling force, 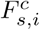 the force due to collision and 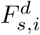 the driving force. Exact descriptions and mathematical equations of the forces implemented here are shown in Supplementary Data 1. We solve the differential equations by iterative Runge-Kutta methods (52) for a total of 150 ms simulation time.

Because the linear connection of the masses produces hard corners and thus, non-natural segmentations, we used the Chaikin’s corner cutting algorithm to produce smooth (more physiological) glottal areas (53). As the model uses six degrees of freedom as initial parameters (Q1 to Q6, similar to (17, 54)), we draw the values for all Qs from a uniform, random distribution within a physiological range (54) to generate a variety of different time-variant segmentation masks. The boundaries were [0.5, 2] for Q1-Q4 (mass and stiffness), [1.0, 4.0] for Q5 (subglottal pressure) and [0.5, 6] for Q6 (collision force). Asymmetric values for Q1-Q4 result in asymmetric oscillations. The first 85 simulated ms were discarded as the model starts in a transient state. The remaining 65 ms were stored for further evaluation. We generated a total of 2500 oscillating models. Models that do not oscillate due to unsuitable initial parameters were discarded.

### Endoscopy data labeling

The BAGLS dataset (36) was labeled manually on each of the 59,250 images using a custom written annotation tool in Python (Supplementary Fig. 8). Locations are stored as (*x, y*) coordinates for posterior and anterior point, respectively. The data is saved in JavaScript Object Notation (JSON) format. Upon publication, we provide the Github repository.

### Classical computer vision methods

All classical computer vision method principles and how they predict the symmetry axis is shown in Supplementary Fig. 1. These methods operate on binary segmentation masks, where pixels belonging to the glottis have a value of 1, otherwise 0.

#### Top and bottom most (TB)

In TB, the top and bottom most point of the segmentation mask in y is selected as posterior and anterior point, respectively. The line connecting the two points is taken as midline (Supplementary Fig. 1a). The following equations generalize the procedure:

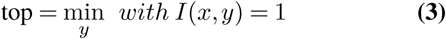

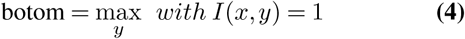

where *I*(*x, y*) is the intensity of the binary segmentation mask with given height (*y*) and width (*x*). If there are multiple px at the same *y* location, the center of mass is used.

#### Linear regression (LR)

Linear regression is a typical operation to fit a line (Eq. S (1)) with a given slope *m* and an intercept *b* to a point cloud by minimizing the residuals in *y* by comparing the true *y* values to the predicted line *ŷ*s (see also Supplementary Fig. 1b):

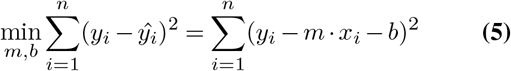

As we are interested in a vertical orientation, we transpose the image, perform linear regression and are using the function inverse to describe the midline. For linear regression, we are using the curve_fit implementation from the Python scipy.optimize library.

#### Orthogonal Distance Regression (ODR)

In ODR, the total least squares of both, *x* and *y*, are minimized, in contrast to linear regression, where only the distance of a given point (*x, y*) in *y* is minimized (55), see also Eq. S (5) and compare Supplementary Fig. 1b and c. This results in minimizing the perpendicular distance *d* from a given point (*x, y*) to the prediction line, which results in point (*x*^∗^, *y*^∗^):

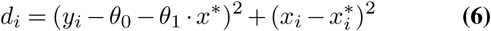

This is computed for all (*x*_*i*_, *y*_*i*_) pairs, resulting in the sum of individual perpendicular distances (*S*), that will be minimized with respect to *θ*_0_, *θ* _1_ and 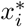:

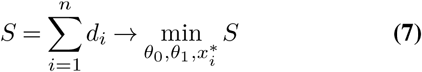

In this study, we use the Python library scipy.odr to interface ODRPACK written in FORTRAN-77 that uses a modified Levenberg-Marquardt algorithm to minimize *S* (55).

#### Principal Component Analysis (PCA)

Here, we utilize an orthogonal linear transformation to translate the image space to a principal component space. As we provide only two dimensions, the first two principal components do fully represent the image data. The first principal component shows the direction of highest variance, i.e. the desired midline vector (Supplementary Fig. 1d).

We first centered our image using the empirical mean for each feature vector, i.e. *x* and *y*, revealing the centroid of our image. We next computed the covariance matrix of *x* and *y*, which was used to find the respective eigenvalues and eigenvectors of the covariance matrix using eigenvalue decomposition. This reveals the first and second principal component. The orientation of the first principal component is used to compute the slope *θ*_1_:

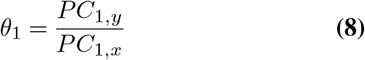

The intercept *θ*_0_ is calculated via the previously revealed centroid of the image.

#### Image Moments

For a binary image with region Ω, the different moments *m*_*pq*_ are calculated from the individual pixels (*x, y*) of Ω ∈ R^2^ as follows:

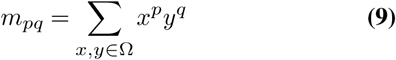

The area |Ω| is defined as the zero-order moment, thus summing the individual pixels:

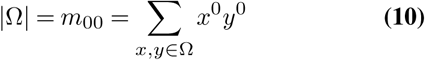

The centroid 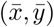 is computed using the center of mass of the region:

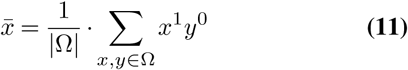

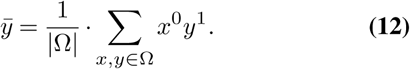

Central Image Moments are by construction translationally, but not rotationally invariant, allowing the estimation of the image orientation (Fig. 3c) and are computed as following:

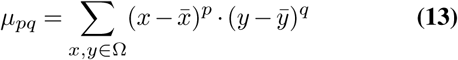

The orientation angle *α*, and thus the slope *θ*_1_, is computed using the following equation:

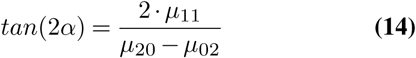

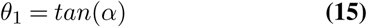

These two features, namely centroid 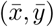 and direction vector, i.e. slope, *θ*_1_ together are predestined to estimate the glottal midline efficiently (Supplementary Fig. 1e).

#### Ellipse Fitting

The glottal area can be approached as an ellipse. By fitting an ellipse to the contour of the segmentation, the major axis would coincide with the midline (Supplementary Fig. 1f).

We are fitting an ellipse of the following form:

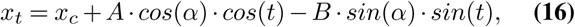

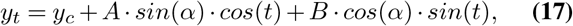

which results in five parameters, (*x*_*c*_, *y*_*c*_) being the centroid of the ellipse, the magnitude of the major and minor axis (*A* and *B*, respectively), as well as the orientation of the ellipse (*α*). As described for the moments, the centroid and the orientation are sufficient to describe the glottal midline. To perform the Ellipse Fitting, we used the contour finding and Ellipse Fitting algorithms in-built in OpenCV. The fitting procedure in OpenCV is implemented according to (56).

### Deep neural networks

All networks were setup in TensorFlow 1.14 with their respective implementation in Keras. All established networks were used from their keras.applications implementation, except EfficientNetB0, where we used the implementation from qubvel. We tested the following loss functions for keypoint prediction (MAE (see Eq. S (18)), MSE (see Eq. S (19)), Huber (see Eq. S (21), as shown in (57, 58)), Log Cosh (see Eq. S (20), as used in (59))) and for semantic segmentation (Dice Loss Eq. S (22) (60)):

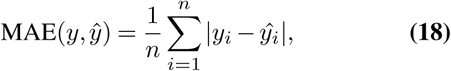

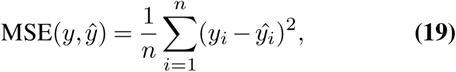

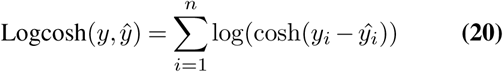

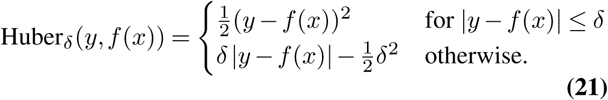

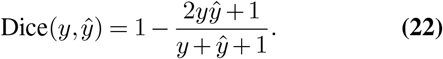

#### Segmentation-based networks

We trained for a maximum of 50 epochs and used eight different starting seeds for crossvalidating training/validation and test-dataset. For each seed, we split the total simulations in 75% for training and 25% for testing. The training set itself was split into 90% training and 10% validation. We used RMSprop as optimizer with 0.9 momentum, a learning rate of 10^−4^ and a learning rate decay of 0.5 · 10^−6^. For evaluation, we used for each architecture the network epoch that performed best on the validation set. MidlineNet is a light-weight variant derived from the U-Net encoder, and features four blocks of two convolutional layers (filter=32, kernel size=3) and a max pooling layer. ReLU was used as activation function. After the four blocks we applied a global average pooling layer and fed this into a dense layer predicting the *x, y* coordinates of the anterior and posterior point.

#### Endoscopy image-based networks

We used the U-Net architecture (35) as basis architecture as implemented previously (10). Additional to the vanilla encoder, we tested the MobileNetV2, the EfficientNetB0 and the Xception encoder with their implementations in Keras. We trained all networks for 100 epochs using either the MAE, the MSE, the Huber or the Logcosh loss for the anterior and posterior point and the Dice loss for the segmentation. The final GlottisNet architecture was trained on the MSE and the Dice loss.

### Evaluation metrics

To evaluate the performance of deep neural networks and the classical computer vision algorithms how well they can predict the glottal midline, we use two metrics: the mean absolute percentage error (MAPE) and the intersection of the union (IoU).

The MAPE metric (Eq. S (23)) defines how close the predicted anterior or posterior point is in respect to the ground-truth values.

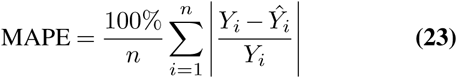

We further use the IoU metric to indicate how much the left and right glottal area provided by the ground-truth midline (i.e. *GT*) overlaps with the left and right glottal area predicted by any algorithm presented here (*P*, Supplementary Fig. 2). We compute the IoU for each side individually and average across both sides to gain the mean IoU (mIoU, Eq. S (25)).

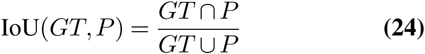

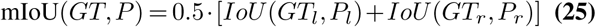

The IoU metric is also used to measure the segmentation performance of the GlottisNet architecture. Here, *GT* is the ground-truth segmentation and *P* the segmentation prediction of the neural network.

## AUTHOR CONTRIBUTIONS

AMK and MD conceived the project. AMK evaluated computer vision algorithms, ported 6MM to Python, generated the model data, annotated the data, designed and trained neural networks and interpreted the data, prepared figures. JZ annotated data and trained neural networks. PG designed 6MM in MATLAB and helped with initial neural networks. MD, AS and SK supervised the project, and acquired funding. AMK wrote the manuscript with the help from all authors.

## COMPETING INTERESTS

The authors declare no competing interests.

## ACKNOWLEDGEMENTS

This work was funded by the BMWi (ZF4010105/BA8) to AMK and MD, by the German Research Foundation DFG (DO1247/8-1, no. 323308998, MD), and a Joachim-Herz-Stiftung Add-On fellowship (AMK).

## Notes

### Competing Interest Statement

The authors have declared no competing interest.

## References

1. Ingo R. Titze and Daniel W. Martin. Principles of voice production. The Journal of the Acoustical Society of America, 104(3):1148–1148, 1998 ISSN 0001-4966. doi: 10.1121/1.424266.s

2. Dimitar D. Deliyski, Robert E. Hillman, and Daryush D. Mehta. Laryngeal high-speed videoendoscopy: Rationale and recommendation for accurate and consistent terminology. Journal of Speech, Language, and Hearing Research: JSLHR, 58(5):1488–1492, 2015 ISSN 1092-4388. doi: 10.1044/2015_JSLHR-S-14-0253.

3. Daryush D Mehta and Robert E Hillman. Current role of stroboscopy in laryngeal imaging. Current opinion in otolaryngology & head and neck surgery, 20(6):429, 2012

4. C. T. Herbst, J. Lohscheller, J. G. vec, N. Henrich, G. Weissengruber, and W. T. Fitch. Glottal opening and closing events investigated by electroglottography and super-high-speed video recordings. Journal of Experimental Biology, 217(6):955–963, 2014 ISSN 0022-0949, 1477-9145. doi: 10.1242/jeb.093203.

5. Hans Larsson, Stellan Herteg??rd, Per-???ke Lindestad, and Britta Hammarberg. Vocal fold vibrations: High-speed imaging, kymography, and acoustic analysis: A prelim-inary report:. The Laryngoscope, 110(12):2117–2122, 2000 ISSN 0023-852X. doi: 10.1097/00005537-200012000-00028.

6. J. Pieter Noordzij and Peak Woo. Glottal area waveform analysis of benign vocal fold lesions before and after surgery. Annals of Otology, Rhinology & Laryngology, 109(5):441–446, 2000 ISSN 0003-4894, 1943-572X. doi: 10.1177/000348940010900501.

7. Ingo R. Titze. Parameterization of the glottal area, glottal flow, and vocal fold contact area. The Journal of the Acoustical Society of America, 75(2):570–580, 1984 ISSN 0001-4966. doi: 10.1121/1.390530.

8. Max-Heinrich Laves, Jens Bicker, Lüder A. Kahrs, and Tobias Ortmaier. A dataset of laryngeal endoscopic images with comparative study on convolution neural network-based semantic segmentation. International Journal of Computer Assisted Radiology and Surgery, 2019 ISSN 1861-6429. doi: 10.1007/s11548-018-01910-0.

9. Mona Kirstin Fehling, Fabian Grosch, Maria Elke Schuster, Bernhard Schick, and Jörg Lohscheller. Fully automatic segmentation of glottis and vocal folds in endoscopic laryngeal high-speed videos using a deep convolutional lstm network. Plos one, 15(2):e0227791, 2020

10. Pablo Gómez, Andreas M Kist, Patrick Schlegel, David A Berry, Dinesh K Chhetri, Stephan Dürr, Matthias Echternach, Aaron M Johnson, Melda Kunduk, Youri Maryin, Anne Schützenberger, Monique Verguts, and Michael Döllinger. Benchmark for automatic glottis segmentation (BAGLS), 2019 type: dataset.

11. J Lohscheller and U Eysholdt. Phonovibrogram visualization of entire vocal fold dynamics. The Laryngoscope, 118(4):753–758, 2008 ISSN 0023852X, 15314995. doi: 10.1097/MLG.0b013e318161f9e1.

12. J. Lohscheller, U. Eysholdt, H. Toy, and M. Dollinger. Phonovibrography: Mapping high-speed movies of vocal fold vibrations into 2-d diagrams for visualizing and analyzing the underlying laryngeal dynamics. IEEE Transactions on Medical Imaging, 27(3):300–309, 2008 ISSN 0278-0062. doi: 10.1109/TMI.2007.903690.

13. Gunnar Björck and Stellan Hertegård. Reliability of computerized measurements of glottal insufficiency. Logopedics Phoniatrics Vocology, 24(3):127–131, 1999

14. Katsuhide Inagi, Aliaa A Khidr, Charles N Ford, Diane M Bless, and Dennis M Heisey. Correlation between vocal functions and glottal measurements in patients with unilateral vocal fold paralysis. The Laryngoscope, 107(6):782–791, 1997

15. J Lohscheller, Hikmet Toy, Frank Rosanowski, Ulrich Eysholdt, and Michael Döllinger. Clinically evaluated procedure for the reconstruction of vocal fold vibrations from endoscopic digital high-speed videos. Medical Image Analysis, 11(4):400–413, 2007 ISSN 1361-8415. doi: 10.1016/j.media.2007.04.005.

16. Rita Patel, Denis Dubrovskiy, and Michael Döllinger. Characterizing vibratory kinematics in children and adults with high-speed digital imaging. Journal of Speech, Language, and Hearing Research, 57(2):S674–S686, 2014

17. K. Ishizaka and J. L. Flanagan. Synthesis of voiced sounds from a two-mass model of the vocal cords. Bell System Technical Journal, 51(6):1233–1268, 1972 ISSN 1538-7305. doi: 10.1002/j.1538-7305.1972.tb02651.x.

18. Raphael Schwarz, Michael Döllinger, Tobias Wurzbacher, Ulrich Eysholdt, and J Lohscheller. Spatio-temporal quantification of vocal fold vibrations using high-speed videoendoscopy and a biomechanical model. The Journal of the Acoustical Society of America, 123(5):2717–2732, 2008 ISSN 0001-4966. doi: 10.1121/1.2902167.

19. Ina Steinecke and Hanspeter Herzel. Bifurcations in an asymmetric vocal-fold model. The Journal of the Acoustical Society of America, 97(3):1874–1884, 1995 ISSN 0001-4966. doi: 10.1121/1.412061.

20. Tobias Wurzbacher, Michael Döllinger, Raphael Schwarz, Ulrich Hoppe, Ulrich Eysholdt, and Jörg Lohscheller. Spatiotemporal classification of vocal fold dynamics by a multimass model comprising time-dependent parameters. The Journal of the Acoustical Society of America, 123(4):2324–2334, 2008

21. Brian A Pickup and Scott L Thomson. Influence of asymmetric stiffness on the structural and aerodynamic response of synthetic vocal fold models. Journal of biomechanics, 42(14): 2219–2225, 2009

22. Patrick Mergell, Hanspeter Herzel, and Ingo R Titze. Irregular vocal-fold vibration—high-speed observation and modeling. The Journal of the Acoustical Society of America, 108(6): 2996–3002, 2000

23. M. Döllinger, U. Hoppe, F. Hettlich, J. Lohscheller, S. Schuberth, and U. Eysholdt. Vibration parameter extraction from endoscopic image series of the vocal folds. IEEE Transactions on Biomedical Engineering, 49(8):773–781, 2002 ISSN 0018-9294. doi: 10.1109/TBME.2002.800755.

24. Michael Doellinger and David A Berry. Visualization and quantification of the medial surface dynamics of an excised human vocal fold during phonation. Journal of Voice, 20(3):401–413, 2006

25. Michael Döllinger, N Tayama, and DA Berry. Empirical eigenfunctions and medial surface dynamics of a human vocal fold. Methods of information in medicine, 44(3):384–391, 2005

26. F. Chaumette. Image moments: a general and useful set of features for visual servoing. IEEE Transactions on Robotics, 20(4):713–723, 2004 ISSN 1552-3098, 1941-0468. doi: 10.1109/TRO.2004.829463.

27. Ming-Kuei Hu. Visual pattern recognition by moment invariants. IRE Transactions on Information Theory, 8(2):179–187, 1962 ISSN 0096-1000. doi: 10.1109/TIT.1962.1057692.

28. Kaiming He, Xiangyu Zhang, Shaoqing Ren, and Jian Sun. Deep residual learning for image recognition. 151203385 [cs], 2015.

29. François Chollet. Xception: Deep learning with depthwise separable convolutions. In Proceedings of the IEEE conference on computer vision and pattern recognition, pages 1251–1258, 2017

30. Christian Szegedy, Vincent Vanhoucke, Sergey Ioffe, Jon Shlens, and Zbigniew Wojna. Rethinking the inception architecture for computer vision. In Proceedings of the IEEE conference on computer vision and pattern recognition, pages 2818–2826, 2016

31. Barret Zoph, Vijay Vasudevan, Jonathon Shlens, and Quoc V. Le. Learning transferable architectures for scalable image recognition. 1707.07012 [cs, stat], 2018

32. Mingxing Tan and Quoc V. Le. EfficientNet: Rethinking model scaling for convolutional neural networks. 1905.11946 [cs, stat], 2019

33. Mark Sandler, Andrew Howard, Menglong Zhu, Andrey Zhmoginov, and Liang-Chieh Chen. Mobilenetv2: Inverted residuals and linear bottlenecks. In Proceedings of the IEEE conference on computer vision and pattern recognition, pages 4510–4520, 2018

34. Karen Simonyan and Andrew Zisserman. Very deep convolutional networks for large-scale image recognition. arXiv preprint 1409.1556, 2014

35. Olaf Ronneberger, Philipp Fischer, and Thomas Brox. U-net: Convolutional networks for biomedical image segmentation. 1505.04597 [cs], 2015

36. Pablo Gómez, Andreas M Kist, Patrick Schlegel, David A Berry, Dinesh K Chhetri, Stephan Dürr, Matthias Echternach, Aaron M Johnson, Stefan Kniesburges, Melda Kunduk, et al. Bagls, a multihospital benchmark for automatic glottis segmentation. Scientific data, 7(1): 1–12, 2020

37. Andrew C. Harvey. Forecasting, structural time series models and the Kalman filter. Cambridge Univ. Press, transf. to dig. print edition, 2009 ISBN 978-0-521-40573-7 978-0-521-32196-9. OCLC: 1014123226.

38. Xingjian Shi, Zhourong Chen, Hao Wang, Dit-Yan Yeung, Wai-kin Wong, and Wang-chun Woo. Convolutional LSTM network: A machine learning approach for precipitation nowcasting. In Proceedings of the 28th International Conference on Neural Information Processing Systems - Volume 1, NIPS’15, pages 802–810. MIT Press, 2015 event-place: Montreal, Canada.

39. Andreas M Kist and Michael Döllinger. Efficient biomedical image segmentation on edget-pus at point of care. IEEE Access, 8:139356–139366, 2020

40. Youri Maryn, Monique Verguts, Hannelore Demarsin, Joost van Dinther, Pablo Gomez, Patrick Schlegel, and Michael Döllinger. Intersegmenter variability in high-speed laryngoscopy-based glottal area waveform measures. The Laryngoscope, 2019 ISSN 1531-4995. doi: 10.1002/lary.28475.

41. Ian Goodfellow, Yoshua Bengio, and Aaron Courville. Deep learning. MIT press, 2016

42. Hoo-Chang Shin, Neil A Tenenholtz, Jameson K Rogers, Christopher G Schwarz, Matthew L Senjem, Jeffrey L Gunter, Katherine P Andriole, and Mark Michalski. Medical image synthesis for data augmentation and anonymization using generative adversarial networks. In International workshop on simulation and synthesis in medical imaging, pages 1–11. Springer, 2018

43. Ian Goodfellow, Jean Pouget-Abadie, Mehdi Mirza, Bing Xu, David Warde-Farley, Sherjil Ozair, Aaron Courville, and Yoshua Bengio. Generative adversarial nets. In Advances in neural information processing systems, pages 2672–2680, 2014

44. Yunchen Pu, Zhe Gan, Ricardo Henao, Xin Yuan, Chunyuan Li, Andrew Stevens, and Lawrence Carin. Variational autoencoder for deep learning of images, labels and captions. In Advances in neural information processing systems, pages 2352–2360, 2016

45. Athanasios Voulodimos, Nikolaos Doulamis, Anastasios Doulamis, and Eftychios Protopapadakis. Deep learning for computer vision: A brief review. Computational intelligence and neuroscience, 2018, 2018

46. Lindsey A Parker, Melda Kunduk, Daniel S Fink, and Andrew McWhorter. Reliability of high-speed videoendoscopic ratings of essential voice tremor and adductor spasmodic dysphonia. Journal of Voice, 33(1):16–26, 2019

47. Rita R Patel, Stephen D Romeo, Jessica Van Beek-King, and Maia N Braden. Endoscopic evaluation of the pediatric larynx. In Multidisciplinary Management of Pediatric Voice and Swallowing Disorders, pages 119–133. Springer, 2020

48. Peter S Popolo and Aaron M Johnson. Relating cepstral peak prominence to cyclical parameters of vocal fold vibration from high-speed videoendoscopy using machine learning: A pilot study. Journal of Voice, 2020

49. Awni Y Hannun, Pranav Rajpurkar, Masoumeh Haghpanahi, Geoffrey H Tison, Codie Bourn, Mintu P Turakhia, and Andrew Y Ng. Cardiologist-level arrhythmia detection and classification in ambulatory electrocardiograms using a deep neural network. Nature medicine, 25(1):65, 2019

50. Sarah Webb. Deep learning for biology. Nature, 554(7693), 2018

51. Travers Ching, Daniel S Himmelstein, Brett K Beaulieu-Jones, Alexandr A Kalinin, Brian T Do, Gregory P Way, Enrico Ferrero, Paul-Michael Agapow, Michael Zietz, Michael M Hoffman, et al. Opportunities and obstacles for deep learning in biology and medicine. Journal of The Royal Society Interface, 15(141):20170387, 2018

52. Ernst Hairer, Michel Roche, and Christian Lubich. The Numerical Solution of Differential-Algebraic Systems by Runge-Kutta Methods, volume 1409 of Lecture Notes in Mathematics. Springer Berlin Heidelberg, 1989 ISBN 978-3-540-51860-0 978-3-540-46832-5. doi: 10.1007/BFb0093947.

53. George Merrill Chaikin. An algorithm for high-speed curve generation. Computer graphics and image processing, 3(4):346–349, 1974 tex.publisher: Elsevier.

54. Pablo Gómez, Anne Schützenberger, Stefan Kniesburges, Christopher Bohr, and Michael Döllinger. Physical parameter estimation from porcine ex vivo vocal fold dynamics in an inverse problem framework. Biomechanics and modeling in mechanobiology, 17(3):777–792, 2018

55. Paul T Boggs and Janet E Rogers. Orthogonal distance regression. Contemporary Mathematics, 112:183–194, 1990

56. Andrew W. Fitzgibbon and Robert B. Fisher. A buyer’s guide to conic fitting. In BMVC, 1995 doi: 10.5244/C.9.51.

57. Trevor Hastie, Robert Tibshirani, and Jerome Friedman. The Elements of Statistical Learning: Data Mining, Inference, and Prediction. Springer Science & Business Media, 2013 ISBN 978-0-387-21606-5. Google-Books-ID: yPfZBwAAQBAJ.

58. Peter J. Huber. Robust estimation of a location parameter. The Annals of Mathematical Statistics, 35(1):73–101, 1964 ISSN 0003-4851. doi: 10.1214/aoms/1177703732.

59. Pengfei Chen, Guangyong Chen, and Shengyu Zhang. Log hyperbolic cosine loss improves variational auto-encoder. ICLR 2019, 2018

60. Fausto Milletari, Nassir Navab, and Seyed-Ahmad Ahmadi. V-net: Fully convolutional neural networks for volumetric medical image segmentation. 1606.04797 [cs], 2016

